# Mitogenome diversity of *Aedes* (*Stegomyia*) *albopictus*: Detection of multiple introduction events in Portugal and potential within-country dispersal

**DOI:** 10.1101/2020.02.12.945741

**Authors:** Líbia Zé-Zé, Vítor Borges, Hugo Costa Osório, Jorge Machado, João Paulo Gomes, Maria João Alves

## Abstract

*Aedes albopictus*, along with *Ae. aegypti*, are key arbovirus vectors that have been expanding their geographic range over the last decades. In 2017, *Ae. albopictus* was detected for the first time at two distinct locations in Portugal. In order to understand how the *Ae. albopictus* populations recently introduced in Portugal are genetically related and which is their likely route of invasion, we performed an integrative cytochrome C oxidase I gene (COI)- and mitogenome-based phylogeographic analysis of mosquitoes samples collected in Portugal in 2017 and 2018 in the context of the global *Ae. albopictus* diversity. COI-based analysis (31 partial sequences obtained from 83 mosquitoes) revealed five haplotypes (1 to 5), with haplotype 1 (which is widely distributed in temperate areas worldwide) being detected in both locations. Haplotypes 2 and 3 were exclusively found in Southern region (Algarve), while haplotype 4 and 5 were only detected in the North of Portugal (Penafiel, Oporto region). Subsequent high discriminatory analyses based on *Ae. albopictus* mitogenome (17 novel sequences) not only confirmed a high degree of genetic variability within and between populations at both geographic locations (compatible with the *Ae. albopictus* mosquito populations circulating in Europe), but also revealed two mitogenome mutational signatures not previously reported at worldwide level. While our results generally sustain the occurrence of multiple introduction events, fine mitogenome sequence inspection further indicates a possible *Ae. albopictus* migration within the country, from the Northern introduction locality to the Southern region. In summary, the observed scenario of high *Ae. albopictus* genetic diversity in Portugal, together with the detection of mosquitoes in successive years since 2017 in Algarve and Penafiel, points that both *Ae. albopictus* populations seem to be already locally establish, as its presence has been reported for three consecutive years, raising the public health awareness for future mosquito-borne diseases outbreaks.

**Author Summary:** In 2017, *Aedes albopictus* was reported for the first time in Portugal at two distinct locations, in the premises of a tyre company in Penafiel, in the North, and nearby a golf course in Algarve, a tourism destination in the southernmost country region. The geographical spread of this species is boosted by larvae and desiccation-resistant eggs transport in aquatic trade goods, as tires and aquatic plants, and adult anthropophilic behavior that favors passive land transportation. In Portugal, especially in the Southern region, temperate climate conditions are adequate for adult mosquitoes survival most of the year. In a way to understand the genetic variability of *Ae. albopictus* populations introduced in Portugal, we analyzed 31 cytochrome C oxidase I gene (COI) partial sequences and 17 mitogenome sequences, integrating them in the context of the global *Ae. albopictus* phylogeographic diversity (i.e., 183 COI and 26 mitogenome sequences previously reported at worldwide level). Although COI haplotype 1 predominated, four additional haplotypes (2 to 5) were detected in Portugal. Subsequent in-depth mitogenome analysis revealed considerable genetic diversity, including not only sequences relating to mitogenomes reported mainly from Italy, Japan and China, but also two novel mitogenome mutational signatures.

Our study indicate that *Ae. albopictus* is locally established in Portugal and intra-country dispersal may have already happened, highlighting the challenges for vector surveillance and control programs aiming at restraining arbovirus disease burden in the future.

## Introduction

*Aedes* (*Stegomya*) *albopictus*, originally described by Skuse in 1894 from India, is one of the most invasive mosquito species that in the last 50 years has successfully colonized most of the tropical and temperate regions worldwide. In the 1970s its expansion was noticed in several islands in the Indian and Pacific Oceans (namely in the Hawaiian Islands) [1] and for the first time in Europe, in Albania, in 1979 [2]. In the 1980s, *Ae. albopictus* was reported in the Americas, becoming established in Harris County, Texas, where it became a dominant vector species in Houston area [3], and in São Paulo, Brazil [4], and in Africa (first recorded in South Africa in 1989) [5]. The geographic range of this species increased dramatically in the 1990s. In Central America, the spread to Mexico also happened in the early 1990s [6, 7] and was confirmed by all countries in 2010 [5].

In Italy, it was detected in 1990, in the port of Genoa where it was introduced in a shipment of used tires from USA [8]. Italy is nowadays considered the most heavily-infested country in Europe, since *Ae. albopictus* become established in most areas of the country (less than 600 m above sea level) and is abundant in many urban areas [9].

Since the introduction in Italy, *Ae. albopictus* has been gradually spreading in Europe, and specially into most of the Mediterranean countries (S1 Fig). More recently, in 2017, this mosquito was reported in Portugal at two distinct locations, in a tyre company located in the North of Portugal, Penafiel (Oporto region) [10], and nearby a golf resort in the South, Algarve region [11]. Since then, its presence has been reported at the same locations continuously and its establishment and dispersal raises concern for autochthonous mosquito-borne disease outbreaks.

The worldwide successful expansion of *Ae. albopictus* has been promoted by unwilled transport of eggs in artificial and natural containers (with several introductions routed by used tires and Lucky bamboo trades), and to its anthropophilic behavior that promotes a close relation with humans and consequent passive transport via private or public ground vehicles [12, 13].

This impressive invasive capacity is undoubtedly associated to this species adaptive plasticity and the significant genetic population-based variation observed [14]. *Aedes albopictus* ability to inhabit temperate regions with relatively cold and dry climates is related to egg diapause which confers cold-hardiness, and is absent in tropical populations of this species, adapted to warm and wet climates [15]. In this sense, the risk of establishment is believed to be related to the origin of the mosquitoes, since mosquito populations with egg diapause of temperate origins are more likely to establish in temperate latitudes [12].

*Aedes albopictus*, beyond being a nuisance species having considerable impact in environmental health and community welfare, is a competent vector species of a wide range of arboviruses and parasites, which raises the most concern in veterinary and public health. The transmission and spread of pathogenic flaviviruses such as Dengue, Zika, West Nile, Yellow fever and Japanese encephalitis viruses, alphaviruses like chikungunya virus, and also bunyaviruses as the La Crosse and Rift Valley fever viruses makes this mosquito a major global public health issue.

Autochthonous transmission of dengue and chikungunya has been reported in Europe related to *Ae. albopictus*, since 2007, when an outbreak of chikungunya with *circa* 330 suspected and confirmed cases occurred in the region of Emilia Romagna in Italy [16, 17]. More recently, chikungunya outbreaks have been reported in France in 2010 [18, 19], 2014 [20], and 2017 [21], and again, in Italy in 2017 [22]. Autochthonous dengue cases caused by dengue serotypes 1 and 2 have also been reported in 2010 in Croatia [23] and France [24], and in France in 2013 [25], 2014 [26] and 2015 [27]. In 2018, 12 cases of autochthonous dengue were confirmed in the EU, six in Spain (five in the region of Murcia and one in Catalonia) and six in France (five cases in Saint Laurent du Var, one case in Montpellier) [28].

In Portugal, a National Vector Surveillance Network—REVIVE (REde de VIgilância de VEctores)— is established since 2008 under the custody of the Portuguese Ministry of Health [29]. Nowadays, the REVIVE network includes the General Directorate of Health (DGS), the five Regional Health Administrations (ARS) (namely Algarve, Alentejo, Lisboa e Vale do Tejo, Centro and Norte), the National Institute of Health Doutor Ricardo Jorge (INSA), and, in the outermost regions, the Institute of Health Administration of Madeira and the Regional Health Directorate of Azores. REVIVE carries out the nationwide surveillance of the most significant hematophagous arthropods in public health (mosquitoes, ticks, and sandflies). Surveillance of mosquito species and screening of field-collected mosquitoes for arboviruses is regularly performed. At airports, ports, storage areas, and specific border regions with Spain, monitoring takes place throughout the year with the commitment of local and regional authorities.

Mitochondrial DNA genes, namely cytochrome oxidase subunit I (COI) and NADH dehydrogenase subunit 5 (ND5), have been largely used to study the genetic relationships of *Ae. albopictus* [30–33] producing significant sequence data from most of the countries where this species has already been recorded. Mitochondrial DNA genes are ideal genetic markers to assess ancestry and demographic changes in populations [34] since their inheritance is uniparental (maternal in *Ae. albopictus*), recombination events are absent, they have high mutation and nucleotide substitution rates and a well-defined effective population size of one-fourth nuclear genes [34–36]. However, some limitations detected in COI and ND5 population studies show that the variation observed may be inadequate to identify and phylogenetically link haplogroups [37–39]. In a way to overcome these constrains, Battaglia et al. [40] studied sequence variation in the entire coding regions of 27 mitogenomes and define five haplogroups in Asia, of which only three (A1a1, A1a2, and A1b) were likely to be related to the worldwide spread of the tiger mosquito.

In order to understand how *Ae. albopictus* populations recently introduced in Portugal are genetically related and which is their likely route of invasion, we performed an integrative COI- and mitogenome-based phylogeographic analysis of mosquitoes collected in Portugal in 2017 and 2018 in the context of the global *Ae. albopictus* diversity.

## Methods

### Mosquito samples and DNA extraction

In order to explore the mitogenome diversity, we analyzed nine *Aedes albopictus* mosquitoes from 12^th^ September to 4^th^ October 2017, and in 11^th^ July 2018 in the premises of a tyre company, located in the metropolitan area of Oporto, municipality of Penafiel, and 17 *Ae. albopictus* females in Loulé municipality, Algarve region, from 12^th^ July 2018 to 4^th^ October 2018 (Table 1). All mosquito samples were collected by the national REVIVE surveillance network at public and private proprieties with the respective accountable/owners knowledge and permission. Samples for DNA extraction were selected individually and in pools (up to six mosquitoes). All these mosquitoes were previously identified at morphological [41, 42] and molecular (Osório et al. [10], and this work for mosquitos collected in 2018) levels. Additionally, 7 and 60 mosquitos, collected in Algarve and Penafiel, respectively, from 26^th^ September to 17^th^ October 2018 were also molecularly identified using COI gene of mitochondrial DNA using primers LCO 1490 and HCO 2198 [43], as previously described [10]. Mosquito samples collected in 2018 were selected for mitogenome sequence using COI haplotype data.

**Table 1.** *Aedes albopictus* samples collected in Portugal.

| Original Designation | COI<br>GenBank ID | mitDNA<br>GenBank ID | Collection<br>Date | Collection<br>Place | Region | Nº Mosq | Mitogenome* | COI<br>Haplotype |
| --- | --- | --- | --- | --- | --- | --- | --- | --- |
| PoMo1076<sup>a</sup> | MF990905 | ND | 04/09/2017 | Penafiel | Oporto | 1 |  | 1 |
| PoMo2599 | MK995303 | MN513352 | 04/10/2017 | Penafiel | Oporto | 1 | New 2 | 1 |
| PoMo2600 | MK995304 | MN513353 | 02/10/2017 | Penafiel | Oporto | 1 | (A1a1a1a) | 1 |
| PoMo2601 | MK995305 | MN513354 | 02/10/2017 | Penafiel | Oporto | 1 | (A1a1a1a) | 1 |
| PoMo2602 | MK995306 | MN513355 | 04/10/2017 | Penafiel | Oporto | 1 | (A1a2) | 1 |
| PoMo2604 | MK995307 | MN513356 | 04/10/2017 | Penafiel | Oporto | 1 | (A1a1a1a) | 1 |
| PoMo2607 | MK995308 | MN513357 | 13/09/2017 | Penafiel | Oporto | 1 | (A1a2a) | 1 |
| PoMo2608 | MK995309 | MN513358 | 12/09/2017 | Penafiel | Oporto | 1 | (A1a1a1a) | 1 |
| PoMo2605/2609/2611<sup>b</sup> | MK995310 | ND | 04/10/2017 | Penafiel | Oporto | 3 |  | 1 |
| PoMoF502 | MK995311 | ND | 11/07/2018 | Penafiel | Oporto | 1 |  | 1 |
| PoMo2708 | MK995312 | MN513359 | 27/09/2018 | Almancil | Algarve | 1 | New 2 | 1 |
| PoMo2711 | MK995313 | MN513361 | 26/09/2018 | Almancil | Algarve | 4 | New 2 | 1 |
| PoMo2713A | MK995314 | ND | 08/10/2018 | Almancil | Algarve | 2 |  | 1 |
| PoMo2724 | MK995315 | ND | 17/10/2018 | Penafiel | Oporto | 6 |  | 1 |
| PoMo2725A | MK995316 | ND | 17/10/2018 | Penafiel | Oporto | 1 |  | 1 |
| PoMo2727 | MK995317 | ND | 04/10/2018 | Almancil | Algarve | 6 |  | 1 |
| PoMoF607<sup>c</sup> | NA | MN513366 | 04/10/2018 | Almancil | Algarve | 1 | New 1 | 2 |
| PoMo2729B | MK995318 | ND | 04/10/2018 | Almancil | Algarve | 3 |  | 1 |
| PoMoF506 | MK995319 | MN513365 | 12/07/2018 | Quarteira | Algarve | 1 | New 1 | 2 |
| PoMo2710 | MK995320 | ND | 26/09/2018 | Almancil | Algarve | 1 |  | 2 |
| PoMo2712A | MK995321 | ND | 08/10/2018 | Almancil | Algarve | 5 |  | 2 |
| PoMo2714 | MK995322 | ND | 04/10/2018 | Almancil | Algarve | 6 |  | 2 |
| PoMo2725B | MK995323 | ND | 04/10/2018 | Almancil | Algarve | 5 |  | 2 |
| PoMo2726 | MK995324 | ND | 04/10/2018 | Almancil | Algarve | 6 |  | 2 |
| PoMo2729A | MK995325 | ND | 04/10/2018 | Almancil | Algarve | 3 |  | 2 |
| PoMo2709 | MK995326 | MN513360 | 27/09/2018 | Almancil | Algarve | 3 | (A1a1a) | 3 |
| PoMo2712B | MK995327 | ND | 08/10/2018 | Almancil | Algarve | 5 |  | 3 |
| PoMo2713B | MK995328 | ND | 08/10/2018 | Almancil | Algarve | 2 |  | 3 |
| PoMo2715 | MK995329 | ND | 04/10/2018 | Almancil | Algarve | 6 | (A1a1a) | 3 |
| PoMoF636<sup>d</sup> | NA | MN513368 | 04/10/2018 | Almancil | Algarve | 1 | (A1a1a) | 3 |
| PoMo2728 | MK995330 | MN513362 | 04/10/2018 | Almancil | Algarve | 6 | (A1a1a) | 3 |
| PoMoF618<sup>e</sup> | NA | MN513367 | 04/10/2018 | Almancil | Algarve | 1 | (A1a1a) | 3 |
| PoMoF505 | MK995331 | MN513364 | 11/07/2018 | Penafiel | Oporto | 1 | (A1a2) | 4 |
| PoMoF503 | MK995332 | MN513363 | 11/07/2018 | Penafiel | Oporto | 1 |  | 5 |

\*Mitogenome designation as defined by Battaglia et al. [40].
<sup>a</sup> [10]
<sup>b</sup>Sequence obtained from 3 mosquitos.
<sup>c</sup>Female mosquito analysed individually from pool PoMo2727.
<sup>d</sup>Female mosquito analysed individually from pool PoMo2715.
<sup>e</sup>Female mosquito analysed individually from pool PoMo2728.
NA – not applicable; ND – not determined.

Mosquitoes were grinded individually or in pools (up to 6 specimens) with a mortar and pestle with liquid nitrogen and 500 µL of minimal essential medium supplied with 10% FBS, streptomycin (0.1 mg/mL) and amphotericin B (1 mg/mL). An aliquot of 300 µL was preserved at −80ºC and the remaining volume was further grinded 300 µL of Lysis Buffer (NUCLISENS® easyMAG, Biomérieux), added to the homogenizer cartridge (Invitrogen) and centrifuged at 12,000g for 2 min to remove cellular debris and reduce lysate viscosity. Total nucleic acid extraction was performed using the prepared lysate suspensions in the automated platform NUCLISENS® easyMAG (Biomérieux).

### Amplicon-based next-generation sequencing

Mitochondrial coding regions (1-14,893 bp) were amplified according to the protocol described by Battaglia et al. [40] by the amplification of two long PCR fragments using primers 274F (5’AGC TAA CTC TTG ATT AGG GGC A3’) and 8875R (5’TGT TGA GGC ACC TGT TTC AG3’) for coding region 1 (8.6 Kb), and 8415F (5’TTA AAG TCG GAG GAG CAG CT3’) and 14717R (5’AAA TTT GTG CCA GCT ACC GC3’) for coding region 2 (6.3 Kb). Long PCRs were carried out in 50 µL reaction mixtures with 1x AccuPrime™ PCR Buffer II (Invitrogen), 1 U of AccuPrime™ *Taq* DNA Polymerase High Fidelity (Invitrogen), 0.2 µM of each primer and 10-50 ng of template DNA. PCR conditions were as follows: denaturation at 94ºC for 2 min, and 35 cycles of 94ºC for 30s, 59ºC for 30 s and 68ºC for 9 min, and final extension at 68ºC for 5 min. Successful amplicons were screened on a 1.5% agarose gel and further purified using Agencourt AMPure XP PCR Purification kit before proceeding to Nextera XT DNA Library Preparation (Illumina), according to manufacture instructions. Libraries were subsequently sequenced (2 x 150 bp or 2 x 200 bp paired-end reads) using a MiSeq (Illumina) equipment.

### Data analysis

Core bioinformatics analyses were conducted using INSaFLU (https://insaflu.insa.pt/), a web-based platform for amplicon-based NGS data analysis [44]. Briefly, the bioinformatics pipeline (detailed in Borges et al. [44]) involved: i) raw NGS reads quality analysis and improvement using FastQC v. 0.11.5 (https://www.bioinformatics.babraham.ac.uk/projects/fastqc) and Trimmomatic v. 0.27 [45] (http://www.usadellab.org/cms/index.php?page=trimmomatic), respectively; ii) reference-based mapping, consensus generation and variant detection using the multisoftware tool Snippy v. 3.2-dev (https://github.com/tseemann/snippy) (the captured sequence of the mitogenome of *Ae. albopictus* strain Rimini isolate 1#Rim1 haplogroup A1a1a1 was used as reference; NCBI accession number KX383916; positions 283-14702); and, iii) alignment of consensus sequences using MAFFT v. 7.313 [46] (https://mafft.cbrc.jp/alignment/software/). Mean depth of coverage *per* sample ranged from ~450x to 1500x. Reads datasets generated during this study are available at the European Nucleotide Archive (Project accession number PRJEB32796). Detailed ENA accession numbers are described in S2 Table.

For the integration of mosquitos circulating in Portugal into the global *Ae. albopictus* genetic diversity, nucleotide consensus sequences of both COI gene and mitogenome were aligned against multiple sequences available at GenBank (183 COI and 26 mitogenome sequences previously reported at worldwide level; S1 and S2 Tables respectively) using MAFFT v. 7.313 [46]. The obtained nucleotide alignments were manually inspected/corrected using MEGA 7.0 [47] (https://www.megasoftware.net/) and further used to build approximately-maximum-likelihood phylogenetic trees applying the double-precision mode of FastTree2 under the General Time-Reversible (GTR) model (1000 bootstraps) [48]. The shared internal regions of the COI gene and mitogenome subjected to comparative genetic analyses in this study correspond to positions 1,511-2,080 and 283-14,653 of the reference mitogenome (Rimini isolate 1; GenBank accession number KX383916), respectively. *Aedes albopictus* metrics of genetic diversity including the number of polymorphic sites, haplotype diversity, and nucleotide diversity for COI and mitogenome sequences, for all determined sequences and for Oporto and Algarve populations were estimated using DnaSP v.5.0 [49].

Phylogenetic data integration and visualization was performed using GrapeTree [50] and Microreact (also used for geospatial data visualization) [51].

## Results

### COI haplotypes diversity in Portugal

Thirty-one mtDNA COI partial sequences from 83 *Ae. albopictus* mosquitoes collected in Portugal were analyzed (GenBank accession numbers MF990905, MK995303-MK995332) representing five haplotypes (i.e. maternal lineages) and a total of four polymorphic sites (Table 1). Estimated nucleotide diversity and haplotype diversity was higher in Algarve than in Oporto population (π= 0.00149, *Hd* =0.6895 vs π= 0.00043, *Hd* =0.2747).

Analysis of partial DNA sequences from the COI gene obtained from mosquitoes collected in Algarve (Southern Portugal) in 2018 confirmed the presence of three haplotypes (1, 2 and 3), and allowed the detection of two new haplotypes (4 and 5) in Penafiel (Northern Portugal) in 2018, besides the haplotype 1 mosquito samples previously collected in this region in 2017 (Table 1 and S2 Fig). Haplotype 1, which is shared in both regions, represents the most common and widely distributed in temperate areas (Fig 1, S2 Fig, Table 1 and S1 Table). The observed diversity detected in Portugal, considering the geospatial haplotype distribution in Europe (Fig 1B; S2 Fig), can be well explained by passive land-transportation from other European countries, especially from the Mediterranean countries.

**Fig 1.**
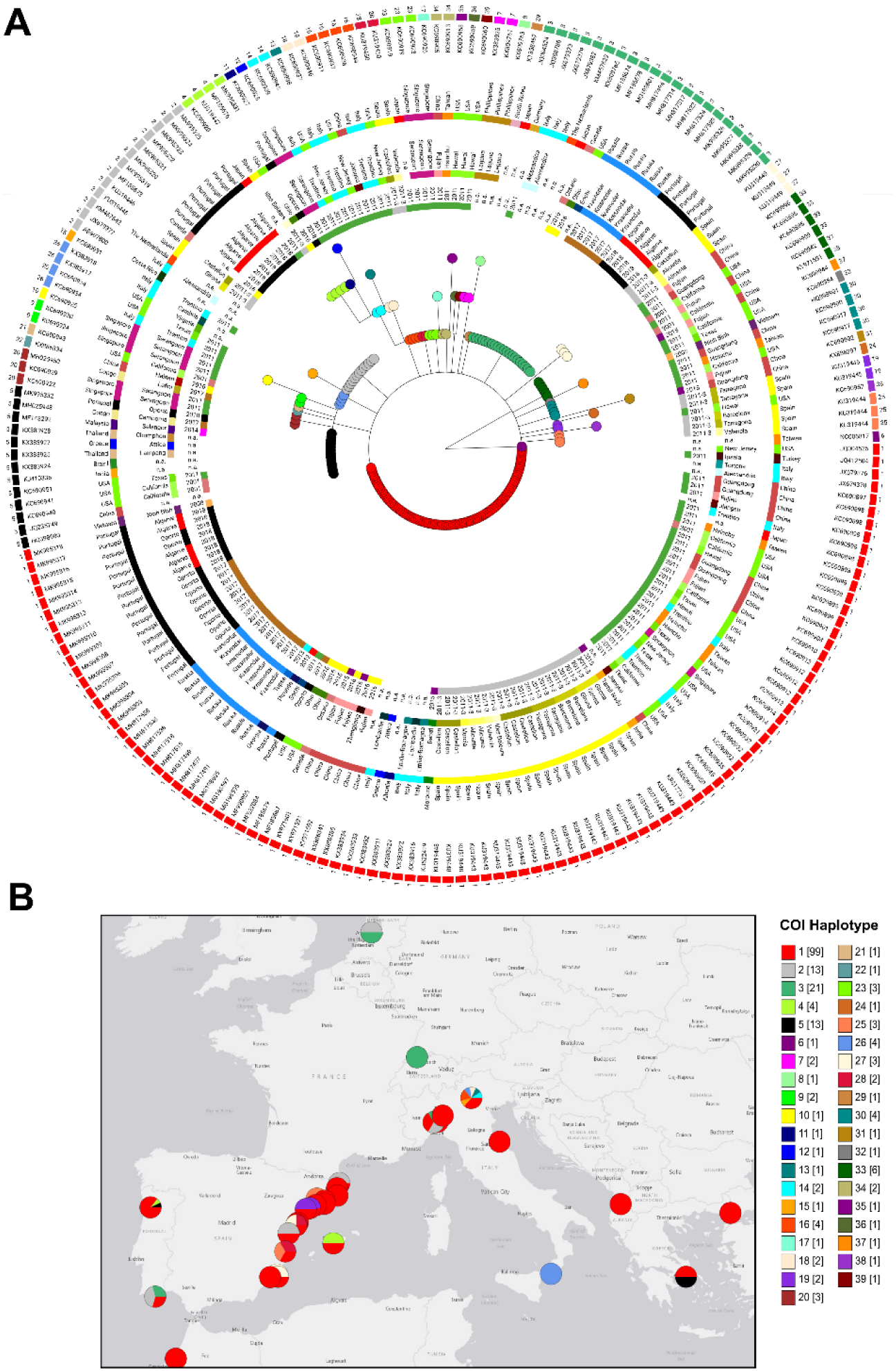
Integration of mosquitos detected in Portugal into the global *A. albopictus* COI-based genetic diversity. **A.** Microreact Visualization of a maximum likelihood phylogenetic tree constructed based on 31 novel COI sequences obtained from mosquito circulating in Portugal plus 182 sequences available at GenBank (S1 Table). The colored external rings (from the outside in) indicate the COI haplotype/GenBank accession numbers, country, country region and year of collection. The tree nodes are colored according with the COI haplotype. For better tree visualization, the highly divergent VN103-9 strain haplotype 40 was excluded from the tree and the NC006817 sequence representative of mitogenome haplogroup A3 / COI haplotype 6 was used as root. **B.** Geospatial mapping of *A. albopictus* detected in the Portugal by COI haplotype in the context of the mosquito distribution in Europe region (plus Morocco). Of note, the geographical placement of circles (colored by haplotype distribution) in the map may not correspond to the exact location where mosquitos were collected (refer to S1 Table for details about the used location) and the circles size does not correlate with number of sequences documented in each location. The internal region of the COI gene under comparison corresponds to positions 1511-2080 of the Rimini isolate 1 reference mitogenome (GenBank accession number KX383916). Phylogenetic and geospatial data were integrated using the freely available platform Microreact [51], with the map presented here being externally created with the open source website https://landlook.usgs.gov/viewer.html.

### Mitogenome-based phylogeography of *Ae. albopictus*

A total of 17 novel mitogenomes coding sequences, 14 almost complete (14,370 – 14,420 bp; GenBank accessions MN513352-MN513359, MN513361, MN513362, MN513364-MN513366 and MN513368) and three partial sequences (GenBank accessions MN513360, MN513363 and MN513367, for PoMo2709, PoMoF503 and PoMoF618, respectively) were determined in this study, representing nine different sequences (Table 1, S3 Table). Using mitogenome sequences, the estimated nucleotide diversity was higher in Algarve (π= 0.00052 vs π= 0.00047), but the haplotype diversity was higher in Oporto *Ae. albopictus* population (*Hd* =0.8056 vs *Hd* =0.7500). Sequence analysis confirms a high level of mitogenome diversity in both locations compatible with the *Ae. albopictus* mosquito populations circulating in Europe (Fig 2, S2 and S3 Tables). Overall, and as previously reported by Battaglia et al. [40] for most of mitogenomes circulating in temperate regions, the Portuguese mitogenome’ sequences grouped within haplogroup A1. Specifically, (i) PoMo2600, PoMo2601, PoMo2604, PoMo2608 mitogenomes from Oporto cluster with mitogenomes from Italy and USA (A1a1a1a, using Battaglia et al. haplogroup designation), (ii) PoMo2728 and PoMoF636 mitogenomes from Algarve clusters with A1a1a mitogenome from Japan (J Wa1), (iii) PoMo2602 and PoMoF505 mitogenomes from Oporto clusters more closely with Chinese Foshan sequence, from a laboratory-maintained strain founded in 1981 from mosquitoes from Southeast China (A1a2), and (iv) PoMo2607 mitogenome from Oporto clusters with Ath2 mitogenome from Greece (A1a2a) (Fig 2, S3 Table). Nevertheless, besides the mitogenome-based divergence reported previously [40], two novel sequence clusters were recovered represented respectively by two sequences, PoMoF506 and PoMoF607 from Algarve, and three sequences, PoMo2599 (2017, Oporto) and PoMo2711 and 2708 (2018, Algarve) (Fig 2). The sequence variation observed in the latest group, enrolling similar sequences observed in the northern region in 2017, and posteriorly in the southern region in 2018, may indicate a possible migration of *Ae. albopictus* within the country, compatible with the observed mutational and temporal profiles (S3 Table).

**Fig 2.**
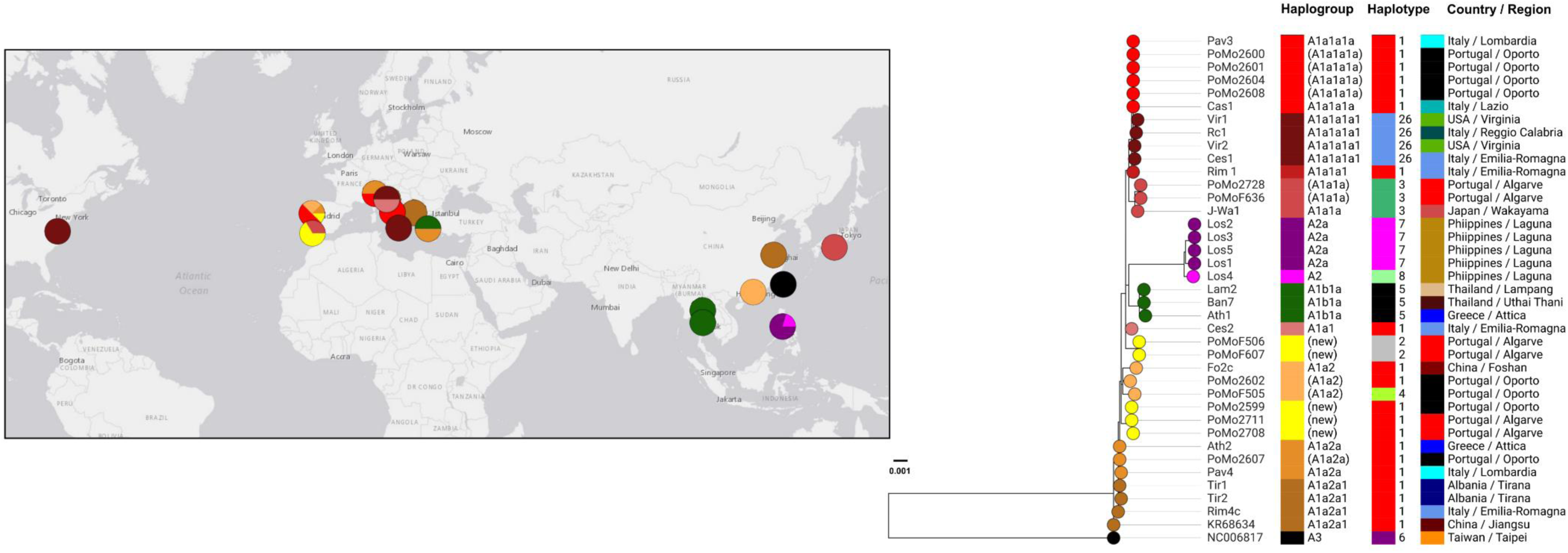
Mitogenome-based phylogeographic analysis of mosquitos detected in Portugal. The figure illustrates the integrative phylogenetic and geospatial analysis of 14 novel mitogenome sequences obtained from mosquito circulating in Portugal plus 25 sequences available in GenBank (S2 Table). The tree nodes are colored according with distinct mitogenome backgrounds (classified as haplogroup, according to Battaglia et al. [40] when possible). Colored blocks (from left to right) indicate the (inferred) haplogroup, COI haplotype and country / region of the 39 mitogenomes under comparison. Of note, inferred haplogroups are indicated within brackets (see details in S3 Table), where divergent sequences that do not present SNP/indel profiles of previously defined haplogroups representative are indicated as potentially “new” (in yellow) haplogroups (including one potentially novel haplogroup detected in Oporto region and another detected in both Oporto and Algarve regions). Of note, the geographical placement of circles (colored by inferred haplogroups) in the map may not correspond to the exact location where mosquitos were collected (refer to S2 Table 2 for details about the applied location). The internal region of the mitogenome sequence under comparison corresponds to positions 283-14653 of the Rimini isolate 1 reference mitogenome (GenBank accession number KX383916). The tree scale reflects the number of substitution *per* site in the 14400 bp alignment. Phylogenetic and geospatial data were integrated using the freely available platform Microreact [51], with the map presented here being externally created with the open source website https://landlook.usgs.gov/viewer.html.

## Discussion and Conclusions

Overall and as expected, the *Aedes albopictus* mosquitos collected in Portugal, in 2017 and 2018, are related to populations involved in the worldwide spread of this species through temperate regions. The genetic diversity observed at both locations, especially by mitogenome sequence analysis, indicates that the main introduction events were distinct and unrelated. However, the determination of a unique cluster of mitogenome sequences in Penafiel, Oporto in 2017 (PoMo2599) and in Algarve in September of 2018 (PoMo2711 and PoMo2708) may indicate migration of *Ae. albopictus* in Portugal from Oporto to Algarve. Algarve region is a common vacation destination for Portuguese population, with increased travel in the summer season. Although in 2019, no additional collections sites for *Ae. albopictus* were, so far, reported by the REVIVE, the geographic spread within the country cannot be excluded. However, further mitogenome surveys enabling the access of sequences from mosquitoes collected in other European countries, namely in Spain are imperative to access a clear picture of this species geographic expansion.

In Penafiel (Northern location, Oporto region), the increased genetic diversity suggests multiple introduction events via tires transport. This hypothesis is supported by the detection of new distinct COI/mitogenome sequences in 2018 comparing with 2017. Despite several international connections by sea and sea ports in Portugal, representing high risk entry points, the pattern of genetic diversity observed in *Ae. albopictus* mosquitoes suggests that the introduction events detected at both sites/regions were promoted by passive land-transportation mainly from other European countries. In Penafiel, at the premises of the tyre company, the introduction event was most probably by immature stages, including eggs and larvae, and in Algarve most likely of adult mosquitoes by passive transport in public or private vehicles. Some tiger mosquitoes mitogenome sequences detected in Penafiel in 2017 (PoMo2600, PoMo2601, PoMo2604 e PoMo2608) are identical to mitogenome sequences detected in Italy (Cassino, region of Lazio [40]; GenBank accession KX383921), indicating this country as the most probable origin of, at least, some of the mosquitoes populations introduced in Portugal. These results are in agreement with the general assumption that Italy is the geographic origin of the recent *Ae. albopictus* spread in Europe [52] and also corroborates the report by Osório et al. [10] of an *Ae. albopictus* Insect Specific Flavivirus (ISF) sequence detected in Penafiel in 2017 similar to ISF sequences detected previously in Italy.

From other perspective, the high genetic diversity perceived in our results prove the coexistence of different genetic sources, thus raising the hypothesis that such population plasticity can be enough to difficult eventual vector control strategies. Additionally, the detection of mosquitoes in successive years since 2017 in Algarve and Penafiel points that both *Ae. albopictus* populations seem to be already locally establish, as its presence has been reported for three consecutive years.

Furthermore, as vector spread is a key risk factor for arbovirus transmission, with arboviruses outbreaks typically occurring 5-15 years after *Ae. aegypti* or *Ae. albopictus* detection [52], the presence of *Ae. albopictus* in Portugal is a major public health threat, raising concern about the introduction of several arboviruses in the continent to the same level as already recognized in Madeira Island. In fact, *Ae. aegypti* present in Madeira since 2005 [53], was associated with the major dengue outbreak reported in Europe in 2012 [54].

Considering the emerging arboviruses (namely Dengue, Zika and Chikungunya viruses) and the rapid movement of viremic travelers, the risk of autochthonous cases in Europe is directly linked to the presence of *Aedes* vectors (especially during the summer season when vectorial capacity is sufficient to sustain transmission) and the number of travelers that arrive viremic increases [55]. A recent work by Massad et al. [55] estimates that Portugal received 71 dengue-viremic air passengers in 2012 (following Germany, France, United Kingdom, Italy and Spain with 167, 150, 148, 124 and 120 estimated dengue-viremic travelers, respectively). This estimate drops to 36 when considering the monthly distribution of the arrivals of Dengue infected passengers from May to October (when vectorial capacity is higher in Portugal). However, in Algarve, which is the southernmost country region, and where more suitable climatic conditions with higher temperatures are observed, the vectorial capacity can be suitable from March-April, until November, when the estimate number of dengue-viremic travelers arriving raises to 64 [55]. Regarding Algarve, the public health concern is thus particularly high, since this region is an important tourism destination in Europe. Moreover, when considering the expected number of dengue viremic air passengers arriving only from Brazil, Portugal is the third European country at high risk, receiving 68 dengue-viremic travelers (closely following France and Italy that are expected to receive 70 and 69 dengue-viremic travelers, respectively). In fact, this work by Massada and co-authors [55] points out the high risk of arbovirus introduction and occurrence of secondary autochthonous cases in Portugal, especially from Brazil.

In conclusion, our work, by providing an unprecedented and detailed picture of the genetic diversity of *Ae. albopictus* detected in Portugal, highlights the importance of surveillance at “hotspots” for mosquitoes introduction, as tires companies, and points of entry (borders, ports and airports), as well as the eventual implementation of vector control methods by the responsible authorities to prevent (new) introductions, establishments and dispersals within the country.

Considering recent prediction studies, habitat suitability for *Ae. albopictus* expansion is expected in Portugal from North to South [52]. In this scenario, the application of vector control measures and the maintenance of national entomological surveillance programs is crucial to effectively lower the risk of future autochthonous arbovirus cases (and outbreaks) in Portugal.

## Acknowledgments

We are grateful to Marco Brustolin and Carles Aranda for proving the collection date and location in Spain of the sampled mosquitoes corresponding to haplotypes H1 to H8 COX1 sequences (Genbank accessions KU319443-KU319450). We are also grateful to REVIVE team for the mosquito collection nationwide.

