## Supplementary figures and images for "Mitogenome diversity of *Aedes* (*Stegomyia*) *albopictus*: Detection of multiple introduction events in Portugal and potential within-country dispersal"

### Supplementary Figure 1

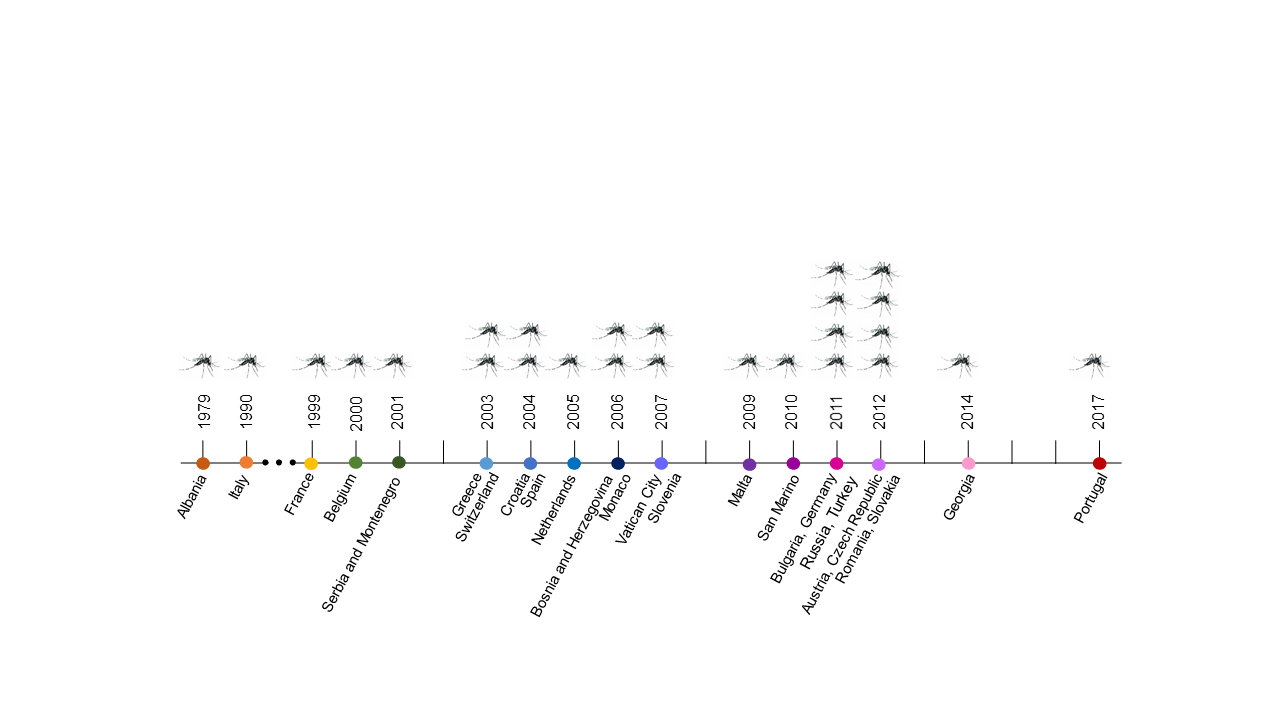

### Supplementary Figure 2

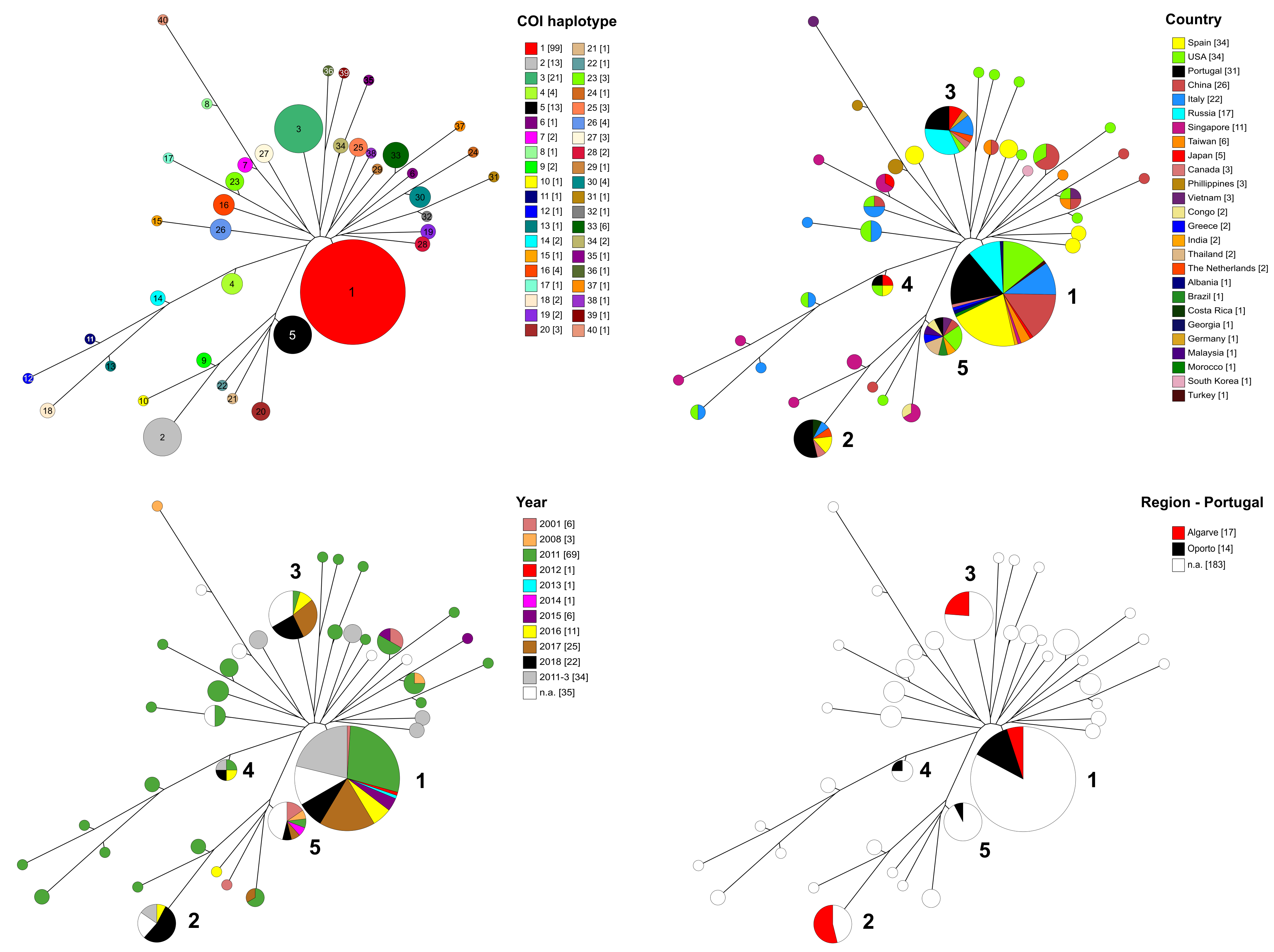
